# Exploiting delayed transitions to sustain semiarid ecosystems after catastrophic shifts

**DOI:** 10.1101/211987

**Authors:** Blai Vidiella,, Josep Sardanyés, Ricard V. Solé

## Abstract

Semiarid ecosystems (including arid, semiarid and dry-subhumid ecosystems) span more than 40% of extant habitats and a similar percentage of human population. As a consequence of global warming, these habitats face future potential shifts towards the desert state characterized by an accelerated loss of diversity and stability leading to collapse. Such possibility has been raised by several mathematical and computational models, along with several early warning signal methods applied to spatial vegetation patterns. Here we show that just after a catastrophic shift has taken place an expected feature is the presence of a ghost, i.e., a delayed extinction associated to the underlying dynamical flows. As a consequence, a system might exhibit for very long times an apparent stationarity hiding in fact an inevitable collapse. Here we explore this problem showing that the ecological ghost is a generic feature of standard models of green-desert transitions including facilitation. If present, the ghost could hide warning signals, since statistical patterns are not be expected to display growing fluctuations over time. We propose and computationally test a novel intervention method based on the restoration of small fractions of desert areas with vegetation as a way to maintain the fragile ecosystem beyond the catastrophic shift caused by a saddle-node bifurcation, taking advantage of the delaying capacity of the ghost just after the bifurcation.

## 1. INTRODUCTION

A major consequence of the habitat degradation processes associated with the Anthropocene is the accelerated loss of habitats and species extinctions that are taking place worldwide (Barnosky *et al.* 2011; Barnosky *et al.* 2012; Hugues 2017). Here importantly, the tempo of this decay is likely to be catastrophic, and evidence from field and experimental data as well as modelling efforts strongly indicate that such decay will end up in collapse (Lenton 2008, Scheffer *et al.* 2001, Scheffer 2009). Ecological shifts are a consequence of the multistable nature of ecosystems, and this is specially relevant in semiarid ecosystems. Indeed, recent field studies in vegetation spatial patterns in drylands have identified multiple ecosystem states (Berdugo *et al.* 2017).

Because of their importance (Maestre *et al.* 2012; 2016) dedicated efforts have been made towards forecasting green-desert catastrophic shifts (Scheffer 2001, Kéfi *et al.* 2007, Scanlon *et al.* 2007, Solé 2007). These *warning signals* would be associated with the presence of increasing fluctuations that are known to diverge close to critical points (Scheffer 2009) as well as to the changes in the distribution of spatial vegetation patterns (Kéfi *et al.* 2014). The definition and quantification of these warning signals has been a very active area over the last decade (Carpenter & Brock 2006; Carpenter *et al.* 2012; Boettiger *et al.* 2013; Scheffer *et al.* 2015). Less attention has been given to another type of phenomenon associated with the behaviour of some nonlinear systems when a parameter *μ* surpasses a critical value *μ*_*c*_. It involves the presence of a special class of long transient or *ghost* (Strogatz1989; Strogatz 2000; Fontich & Sardanyés 2008, Duarte *et al.* 2012). Over this transient, the system will appear stationary, thus creating the false impression that nothing wrong is happening, despite the inevitable ending. In particular, it has been shown that ghosts follow a universal scaling law (Strogatz 2000): the length of transients, 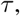 scale following an inverse square-root law, given, in a general form, by

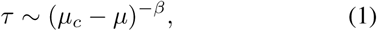

where *β* = 1/2. This scaling law has been identified in physical systems (Strogatz & Westervelt 1989; Trickey & Virgin 1998), in metapopulation mathematical models for autocatalytic species (Sardanyés & Fontich 2010), in low-dimension (Sardanyés & Solé 2006; Sardanyés & Solé 2007) and high-dimensional replicator models exhibiting cooperation (Silvestre & Fontanari 2008; Guillamon *et al.* 2015), as well as in single-species discrete models with Allee effects (Duarte *et al.* 2012).

If ghosts appear in green-desert transitions, the take home message is simple but leads to a grim picture: a given ecosystem might be on its way towards collapse despite the apparent stability. In this paper we show that ghosts are indeed expected in the dynamics of semiarid ecosystem models. However, a positive message is that the ghost can be exploited to ensure ecosystems’ persistence. We introduce a simple intervention method based on the recovery of small fractions of desert areas with vegetation as a way to keep drylands dynamics near the phase space regions where delays occur, thus allowing the ecosystem to remain stable once the bifurcation towards desertification has taken place.

## 2. MATHEMATICAL MODEL

The starting point of our study will be the model studied by Kéfi and co-workers as a coarse grained approximation to semiarid ecosystems (Kéfi *et al.* 2007). In this context, we will not explicitly take into account spatial structure (Rietkerk et al 2002; Meron 2012), stochastic fluctuations (Ridolfi et al, 2011) nor an explicit coupling with soil moisture (Rodriguez-Iturbe et al 1999; D’Odorico et al 2007) or some sources of heterogeneity (Schneider and Kéfi 2016), which can play a relevant role. Here we aim at understanding the properties and management of long transients close to the green-desert shift. The presence of very long transients can force us to redefine the concept of steady state (Kaneko 1990; Hastings 2001; 2004) and provide essential clues to the persistence and organization of populations over long time scales.

The original model (Kéfi et al 2007) considers transitions between vegetated sites (*V*), non-vegetated but fertile sites (*S*), and destroyed sites (*D*) in a given spatial domain (modelled by means of asynchronous cellular automata). The transition probabilities (*ω*) between these states are given by:

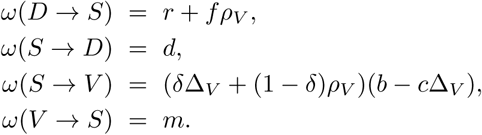

Because of the nature of semiarid ecosystems, the transition probabilities are not only driven by the nearest vegetated neighbours (*ρ_v_*), but also by non-local processes that depend on the fraction of global vegetation (Δ_*v*_). Here, *r* is the spontaneous regeneration rate of a site due to e.g., arrival of seeds by the wind. The parameters *f* and *d* stand for the facilitation and the soil degradation rates, respectively. Regarding the spreading of the vegetation, the dispersion of the seeds are modeled by the constant *δ* (ratio of vegetation due to the whole vegetated sites), the sprout of the seeds *b* and the seed degradation *c*. Finally, the rate of vegetation decay is given by a probability *m* (associated with death or grazing).

The microscopic transition rules provide the rates of growth and death that can be incorporated into a continuous (mean-field) modeling framework. The resulting equations are expressed in terms of balances between different processes:

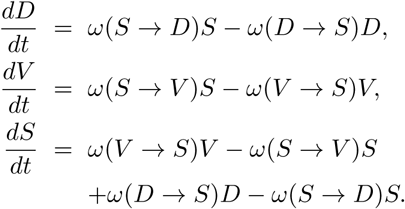

Ignoring spatial correlations we have Δ_*V*_ = *ρ*_*V*_ = *V*. This assumption lead us to the following system of differential equations:

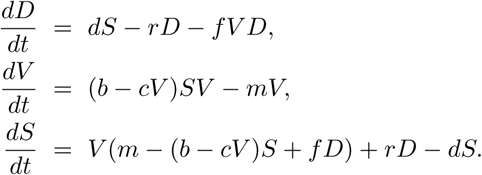

The state variables of the system above correspond to the ratio of the area that is occupied by each different state, with *D+V*+*S* = 1. Thus we can reduce the three-variable system to a two-variable one by means of the linear relation *D* = 1 − (*S + V*). The two-variable system is then:

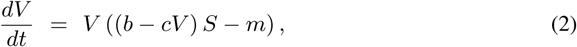

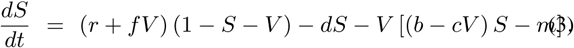

Given that we are modeling the whole semiarid ecosystem, we assume that no income (organic matter or seeds) from outside can occur. Hence, it is assumed that no spontaneous recovery is possible from external sources i.e., *r* = 0.

## 3. RESULTS

### Catastrophic shifts in semiarid ecosystems

The model (2)-(3) displays bistability: for low values of the soil degradation rates, *d*, two possible stable states are present. Actually, this system has three fixed points. The first fixed point is (*V*^*^ = 0, *S*^*^ = 0), which corresponds to the extinction of the vegetation and the destruction of the fertile soil, with a dominance of the desert state. The extinction state is locally stable (see Section S1). The system displays an abrupt transition. This catastrophic shift leads to an *absorbing* state, i.e. once *d* > *d*_*c*_ the system falls into the desert state, with no way of moving away. These results are summarized in Fig. 1, where the fraction of vegetated habitat is plotted against *d.* The increase in *d* involves a monotonous decrease of vegetation. However, at a given critical value *d*_*c*_ an abrupt extinction takes place. Notice that in the bifurcation diagram two fixed points (one stable and one unstable) approach each other as *d* is increased. Specifically, under the parameter values used in our analyses (i.e., *b* = 0.3, *c* = 0.15, *m* = 0.1, and *f* = 0.9) the bifurcation value is *d_c_* ≈ 0.21803399 … (see Section S2 for the description of the computation of this critical value with high accuracy).

**FIG. 1:**
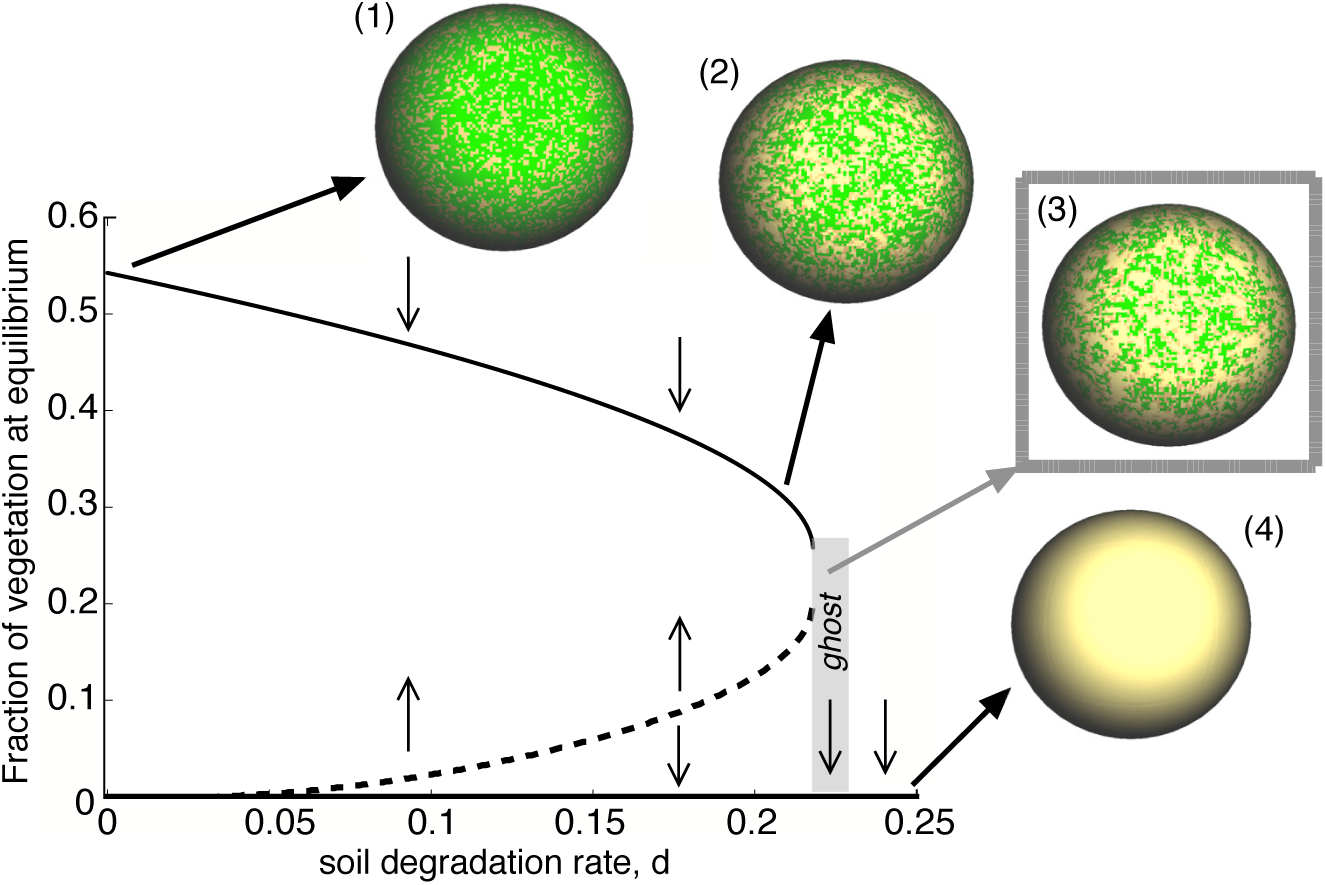
Catastrophic transition found in the model by Kéfi *et al.* (2007). The model considers three variables: vegetated area (*V*), non-vegetated area with fertile soil (*S*), and desert area (*D*). The diagram displays *V* against soil degradation (*d*) at equilibrium using *b* = 0.3, *c* = 0.15, *m* = 0.1, and *f* = 0.9. The upper branch (solid line) and the lower branch (dashed line) are the stable and unstable fixed points. The spheres are spatial snapshots at *t* = 100 using *d* = 0.001 (1); *d* = 0.21 (2); *d* = 0.22 (3); and *d* = 0.25 (4).

The phenomenon that we are investigating here takes place right after the bifurcation. When *d* ≳ *d*_*c*_ the system has a single steady state: the desert state. However, for *d* close enough to *d*_*c*_ the time to extinction *T*_*e*_ rapidly diverges. This means that the system can appear resilient but after a long time it will suddenly collapse. This phenomenon is also depicted in Fig. 1 by means of the representation of the ecosystem in its spatial version. The snapshots in Fig. 1 display the vegetation pattern using four values of *d*. For case (1) the ecosystem is fully vegetated, while for cases (2) and (3) the faction of vegetation is intermediate. However, case (3) is obtained with a value of *d* once the bifurcation has occurred, but the system remains resilient for long time. However, after a very long time, the system eventually decays into (4) i.e., in a completely desert state.

### Dynamics near the desert transition

The detailed dynamics under the values of *d* used in Fig. 1 are displayed in Fig. 2 by means of trajectories in the ( *V*, *S*) space (confined on a triangle i.e., simplex, since *V* + *S* + *D* = 1). When *d* = 0.001 the attractor involving coexistence of states *V* and *S* is close to the side of the simplex, and the basin of attraction of the fixed point (*V* = 0, *S* = 0) is extremely small (see the time series in Fig. 2.1 and snapshot 1 in Fig. 1). As *d* is increased the basin of attraction of the fixed point (0, 0) (shown in red) becomes larger, since the stable node of *V* and *S* coexistence has entered into the simplex approaching to the saddle. For this case, the system is less vegetated (see sphere 2 of Fig. 1.). Notice that in the zoom of Fig. 2.2 the thick trajectory rapidly goes to the stable manifold (SM) of the fixed point and then it slowly approaches to it.

**FIG. 2:**
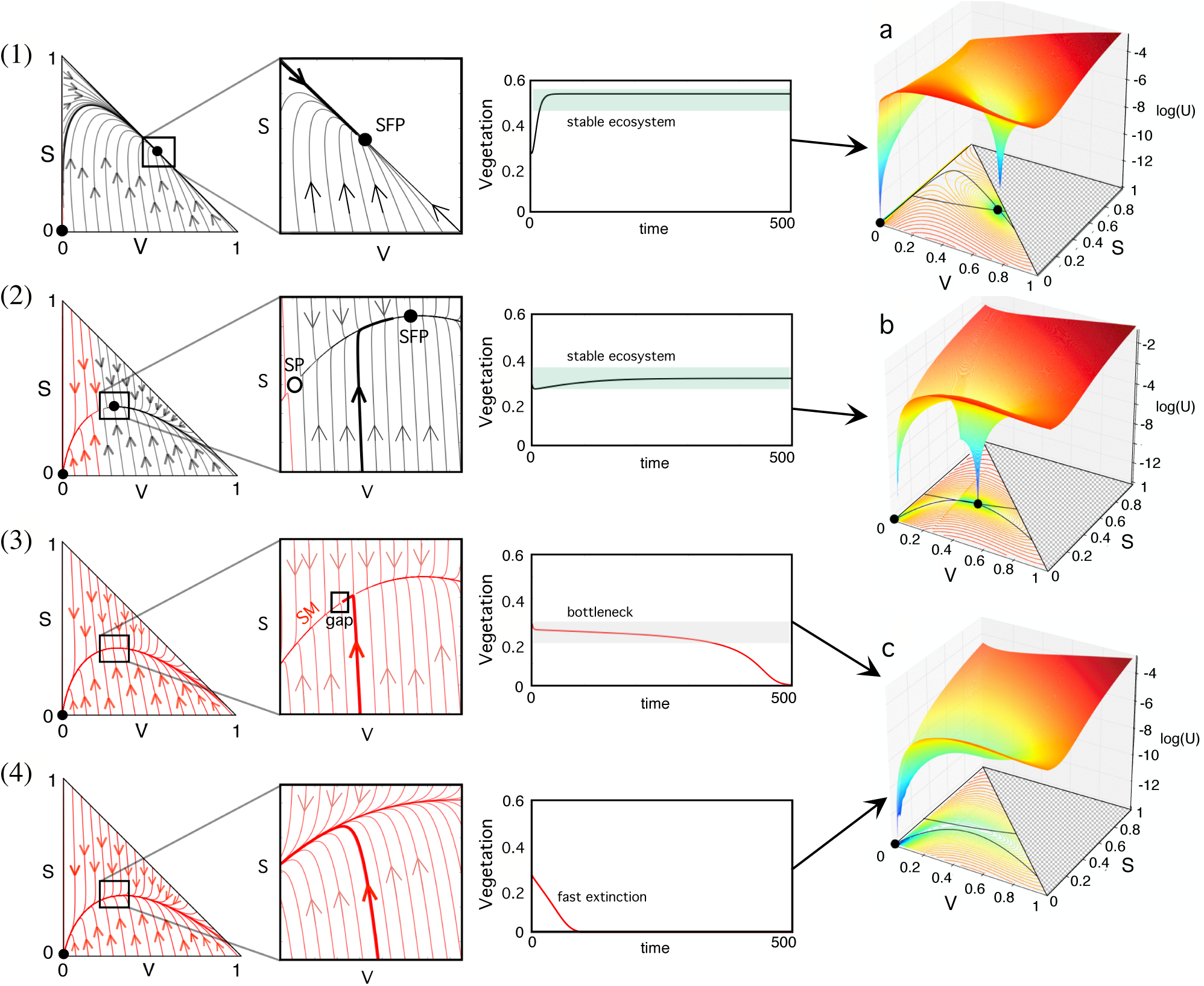
Dynamics at increasing the degradation rate of fertile soil, *d,* with: (1) *d* = 0.001; (2) *d* = 0.21; (3) *d* = 0.22; and (4) *d* = 0.3 (these correspond to the parameters used in Fig. 1). Examples (1) and (2) are computed for *d* < *d_c_*, where the system is bistable (SFP: stable fixed point (black circles); SP: saddle point (white circle)). As *d* is increased (2) the basin of attraction for the desert state grows (illustrated with red orbits). The time series for both cases show persistence of the vegetation. In panels (3) and (4) the critical point *d*_*c*_ has been overcome. While close to *d*_*c*_ (3), the trajectories undergo a slow passage once they reach the stable manifold (SM). The time series displays extinction after a long time. When the value of *d* is further increased extinction occurs rapidly (4). The thick trajectories in the zooms and in the time series in all panels are obtained with the initial condition *V*(0) = 0.28 and *S*(0) = 0.2. Panels (a-c) at the right column: Quasi-potential landscapes before the saddle-node bifurcation (a); at the bifurcation value (b); and once the saddle-node bifurcation has taken place (c). Specifically we use (a) *d* = 0.001, (b) *d* = 0.2, and (c) *d* = 0.3; the other parameters are the same used in the Fig.1. The nullclines (computed at SI.2) for each panel are displayed on the *V-S* plane. The color lines on the plane *V-S* are the isoclines indicating the value of the quasi-potential function. The solid circles are stable fixed points.

The two previous scenarios involved a stable *V* state, provided that *V*(0) and *S*(0) are large enough. However, a more dangerous scenario arises when *d* is slightly above its critical value. This case is shown in Fig. 2.3 using *d* = 0.22. Here notice that all of the trajectories within the simplex travel towards the desert fixed point (0, 0). However, a so-called ghost (or saddle-node remnant) appears in the simplex (close to the place where the stable node and the saddle collided) and it sucks the flows, which undergo a very slow passage through this bottleneck before reaching the point (0, 0). The zoom of Fig. 2.3 reveals that the trajectories rapidly travel to the stable manifold but they undergo a very long passage once they reach this manifold. Notice that for the initial condition used with the thick line, the trajectory flows vertically to the stable manifold and then it remains a very long time before reaching the point (0, 0). A detailed view of the zoom reveals that, for this particular initial condition, the thick trajectory is still flowing in the saddle remnant at *t* = 100 (notice the small gap at the left of the trajectory on the manifold). This phenomenon is displayed in the time series of Fig. 2.3 where a bottleneck is experienced by the trajectory.

The observation of the ecosystem in this scenario over a long transient indicates a seemingly stable state. However, the desert state will be achieved in the long term, and this desertification becomes extremely sharp (see the time series of Fig. 2.3). Here, the observation of the ecosystem at a given time point would result in a seemingly stable system with vegetation (as illustrated with the sphere 3 in Fig. 1). However, as mentioned, after a very long time the system would rapidly collapse towards the desert state. Finally, once *d* grows further away from the critical value *d*_*c*_ transients to extinction are faster. Notice that the same initial condition (thick red trajectory) used in Fig. 2.3 and that did not cover the stable manifold (see zoom in Fig. 2.3) rapidly approaches to this manifold flowing along the entire region covered by the zoom in Fig. 2.4. As a result, the extinction dynamics becomes very fast (see time series in Fig. 2.4).

Another way to characterize the changes in the phase space when tuning the model parameters is given by the so-called potential functions or quasi-potentials (Bhattacharya *et al.* 2011). They provide information on the stability of the fixed points: stable and unstable fixed points appear as valleys and mountains in this functions, respectively. The quasi-potential for Eqs. (2)-(3) in its general form reads

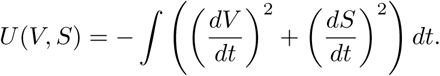

The quasi-potential, computed numerically (see section S3) and displayed at the right hand side of Fig. 2, indicates bistability (for *d* < *d*_*c*_) with two minima placed at the same positions where the nullclines cross. Whereas when there is a unique stable fixed point (for *d* > *d*_*c*_) the quasi-potential function has a single minimum (Fig. 2c). Since the potential function is the integral of the field of Eqs. (2)-(3), the behaviour of the quasi-potential function (Fig. 2) is equivalent to the flows of the phase planes under the same parameter values.

### Transients towards full desert ecosystem

The dependence of the times to extinction, *T*_*e*_, on *d* is displayed in Fig. 3. Figure 3a shows a possible scenario that could have dramatic consequences. If *d* is below its critical value, the ecosystem achieves a stable state. For example, using *d* = 0.217 the fraction of habitat vegetated is about 27% (dashed line in Fig. 3a). For this value of *d* the ecosystem is resilient. However, if *d* is slightly increased (*d* = 0.218034) the system achieves a seemingly stable fraction of vegetated habitat (of about 25%), which actually still holds at *t* ≈ 3.5 × 10^5^. However, since the bifurcation has already occurred, the system rapidly collapses after this extremely long delay at *t* = 352419. The same behavior is found at *d* = 0.21804 and *d* = 0.218055, although here the times to extinction are shorter. Indeed, the dependence on the extinction times on the distance of the bifurcation parameter (*d*) from the bifurcation value (*d_c_*) is shown to follow the inverse square-root scaling law, in agreement with previous works on delayed transitions tied to saddle-node bifurcations (Strogatz & Westervelt 1989; Strogatz 2000; Solé & Sardanyés 2006; Fontich & Sardanyés 2008; Duarte *et al.* 2012). Here *T*_*e*_ is plotted against the distance to the bifurcation value using Θ = *d* – *d*_*c,*_ and the power-law dependence 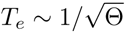 is found numerically.

**FIG. 3:**
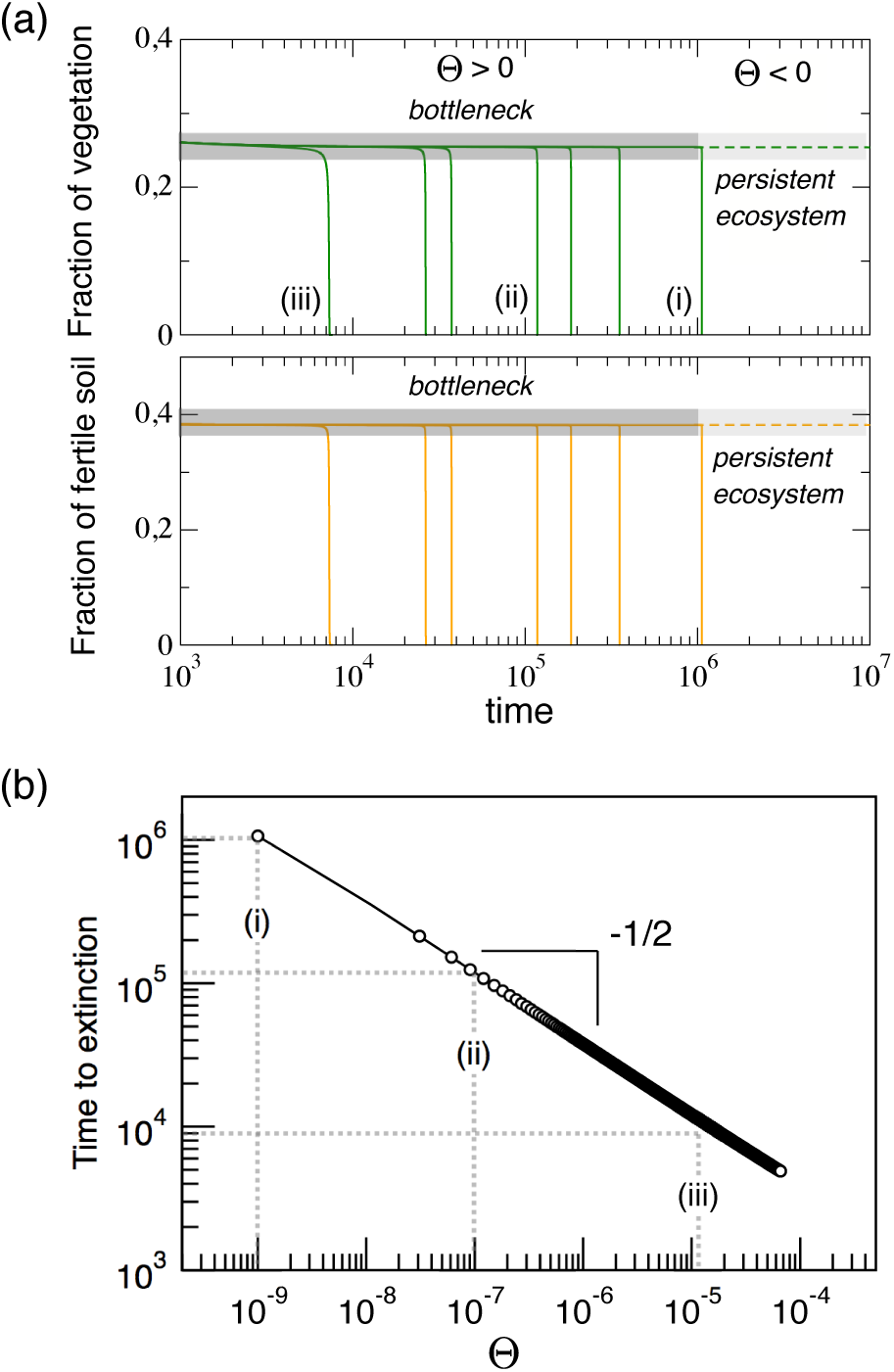
Time to extinction near bifurcation threshold, Θ = *d — d*_*c*_, *d* and *d_c_* being the degradation rate of fertile and the bifurcation value, respectively. (a) Ecosystem persistence below bifurcation threshold (dashed lines, *d* = 0.127 with Θ < 0) and dynamics towards desertification above bifurcation threshold (solid lines with Θ > 0). The time series indicated with (i)-(iii) have been obtained from the values of Θ displayed in panel (b). Notice that in both panels we used the same parameter values used in Fig. 1, except for *d*. (b) Inverse square-root scaling law dependence on extinction transients as the parameter *d* increases above its bifurcation threshold (shown in log-log scale).

The delaying capacities of the ghost are known to depend on the initial conditions (Fontich & Sardanyés 2010). Usually, for initial conditions large enough the ghost captures the orbits and the delays appear. This phenomenon is illustrated in Fig. S3, which displays these times for values of *d* close to the bifurcation computed at different regions of the simplex (by using a regular grid of 10^3^ × 10^3^ different initial conditions). Here, two well-differentiated regions are found. One region where times are extremely long (*T*_*e*_ ≈ 5 × 10^5^ in Fig. S3a) is identified at the right-hand side of the place where the saddle and the node collided (white area). The other region, indicated in black (Fig. S3a), shows those initial conditions for which the orbits go to the desert state very rapidly. Indeed, extremely close to the bifurcation value, these two regions indicate initial conditions spending extremely long transients or initial conditions for which extinction is almost immediate. Once the bifurcation value is further increased times reduce substantially. For instance, Figs. S3b and S3c also display the two regions with long and fast transients. In both cases, long transients are about 1.5 orders of magnitude below the times of Fig. S3a. Note that in Fig. S3 we display time series for values at the right side of the simplex (upper panels with long transients) as well as to the left, where transients are much shorter.

### Control for avoiding catastrophic shifts

The nature of the ghost behavior might provide a way of prolonging the time towards the desert state by using external interventions. Indeed, the maintenance of ecosystems’ stability once the bifurcation has occurred shall be possible and depend on the magnitude of the interventions and their frequency of application. In the case of semiarid ecosystems, the interventions we propose consist on the replantation of vegetation in small fraction of desert areas within the ecosystem. The parameters we consider for the interventions are the fraction of area replanted (expressed as an increased in the fraction of vegetation, Δ*V*) and the frequency (hereafter *freq*) of such a replantation. Notice that the time lapsed during the interventions is assumed to be much smaller than the timescale of the ecosystem dynamics. The intervention strategies that one can consider range from very rare (low frequency) and big areas replanted to small, but very frequent, re-plantations. The effect of different values for both parameters Δ*V* and *freq* could involve three different asymptotic behaviors: (i) ecosystem collapse, (ii) increase of the time delay towards desertification, or (iii) ecosystem preservation.

The response of the ecosystem to a given intervention strategy (in terms of frequency and Δ*V*) depends on the ecosystem condition. In particular, it depends on the increase of the soil degradation rates above the bifurcation value. In order to evaluate and quantify the response of the ecosystem we have numerically simulated the external interventions using a modified 4-th order Runge-Kutta method incorporating the external increase of vegetated area, performed at a given frequency (see section S4 for details on the implemented algorithm, written in pseudo-code form). Figure 4a-c illustrates the intervention processes and the dynamics tied to this process. An oscillatory behavior obtained through the interventions is displayed in the phase portraits (*V, S*) (see Fig. 4a), where the trajectories are constantly re-injected close to the ghost. The effect of the intervention is to preserve (or return) the system into the region where the ghost has a strong delaying effect. The system is thus forced to ”oscillate” because of the interventions (see also Fig. S4). Indeed, this is a slow-fast oscillator because the intervention is instantaneous but then the trajectory enters into the slow passage of the ghost. When the pulses fail in returning the trajectory towards the regions influenced by the ghost the ecosystem collapses via damped oscillations towards the desert state. On the other hand, when trajectories are effectively sent to the ghost, they achieve a stationary cyclic behavior, keeping the system into the vegetated state indefinitely.

**FIG. 4:**
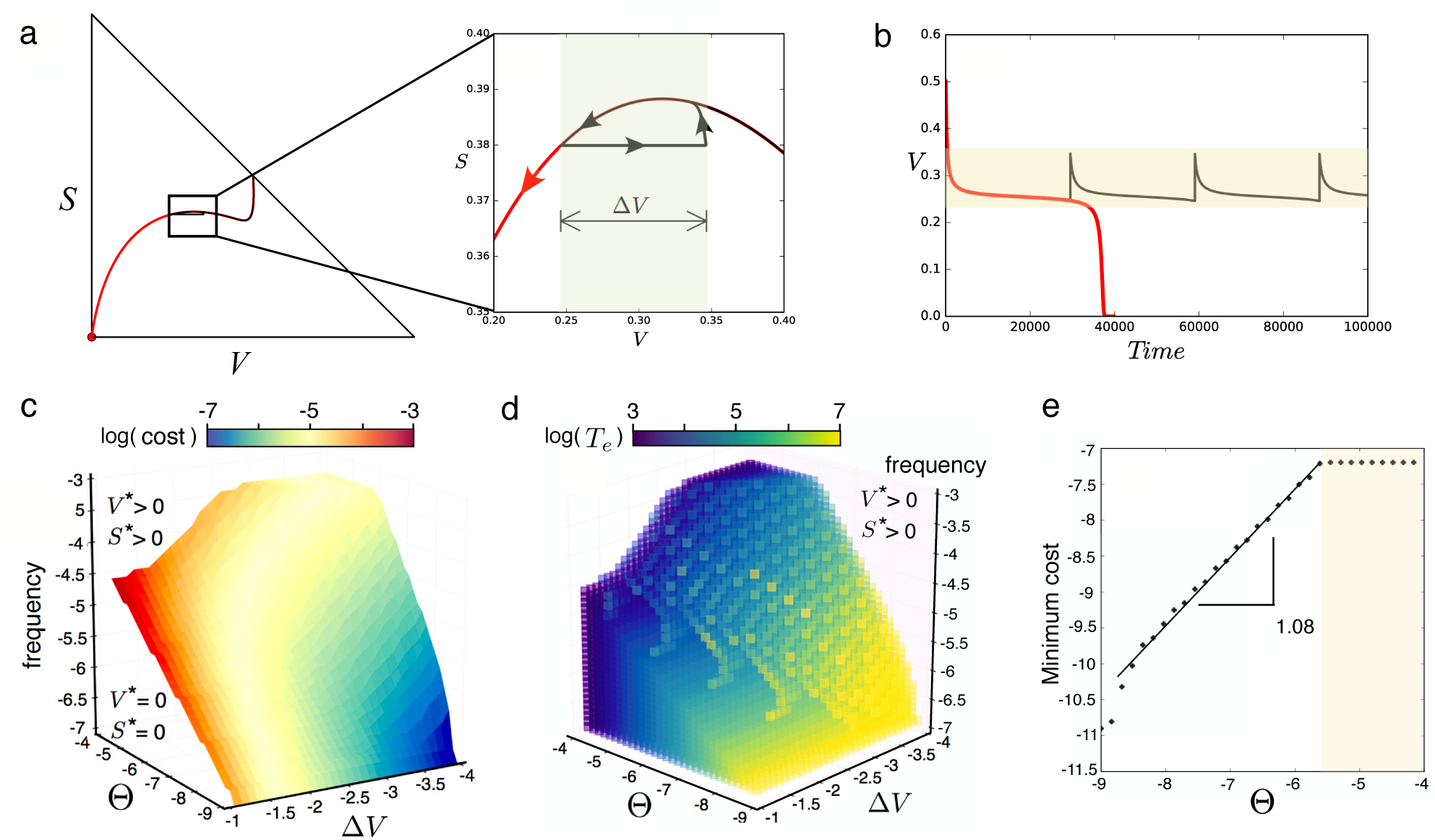
(a-b) Dynamics tied to the interventions. (a) Phase portrait with a trajectory (red) that achieves extinction after suffering a delay of *t* ≈ 4 } 10^4^. Overlapped we display the dynamics using the same initial condition but applying the intervention (black obirt): here using Δ*V* = 0.1 and *freq* = 1/(3 } 10^4^).The enlarged view displays the trajectories during the interventions, which return the orbits near the ghost, giving place to an externally-forced oscillatory-like behavior (black orbit). The red orbit displays the ecosystem extinction without the intervention. (b) Time series for both orbits displayed in (a). The parameters for this calculations are the ones used in the previous figures. (c) Limit between extinction and persistence as a function of the frequency and Δ*V* when increasing *d* above the bifurcation threshold Θ = *d* − *d*_*c*_. (d) Time to ecosystem extinction towards desert in the same space as in (c) i.e., extinction times below the surface displayed in (c). (e) Minimum cost to preserve vegetation at increasing Θ. The straight line shows a power-law fit with exponent 1.08 (with correlation coefficient 0.9965, see also Fig. S5 for further examples). Note that the axes in panels (c-e) are in log-log scale.

In order to quantify how optimal are the interventions we have to take into account that they depend on two parameters (*freq* and Δ*V*). For this reason, we obtain a surface that optimizes the intervention parameters for a given Θ, given by a Pareto front (Fig. 4c). However, to find an optimal intervention strategy for the persistence of a given semiarid ecosystem, it is necessary to design a strategy that optimizes both parameters at the same time. In our case, the optimality of an intervention is evaluated with a simple cost function, defined as the fraction of desert areas replanted by unit time (*plants per area × time*^−1^) that is needed to sustain the system avoiding desertification, with:

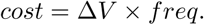

Here the cost is assumed to increase if the fraction of re-planted area and/or the frequency of re-plantation increase. The changes in the cost at increasing the degradation rate are shown in the Fig. 4e. Notice that the relation between the increase of degradation (Θ) and their optimal cost of intervention is almost linear (1.08) in logarithmic scale. The slope of the increase in the optimal cost is twice the decrease rate of the extinction time. This slope depends on the frequency of the interventions and on the are re-planted. In the case of fixing the desired area to be re-planted in each intervention, the optimal frequency will be the one corresponding to the Pareto front surface (Fig. S5a). For bigger amplitudes of the interventions, the increase in the cost function is lower (and the optimal intervention frequency). The slope goes from 0.5 for large interventions (Δ*V*=0.1) to 1.08 for very small re-plantations (Δ*V* = 10^−4^), see Fig. S5b.

As previously mentioned, another possible scenario may be found depending on the magnitude and frequency of the interventions, which could not be enough to achieve the preservation of the ecosystem, but increase the extinction times (see Fig. S6). Figure S6a displays the increase in the extinction times with respect to the non-interventions case in the space (Δ*V*, *freq*) as a function of Θ. A particular example of this delayed transition is shown in Fig. S6b, in which the orbits can be re-injected multiple times until it escapes from the influence of the ghost towards ecosystems’ collapse. Fig S6c displays several cuts done in the three-dimensional space displayed in Fig. S6a. Generically, this kind of interventions are less costly than the ones ensuring full ecosystems’ preservation, and they could be enough to save time since a bigger intervention or ecosystem change can be performed. In the case of semiarid ecosystems, the livestock policy or an active change of the ecosystem (i.e. increase of the humidity) are changes that may return the system to a lower degradation soil condition.

## 4. DISCUSSION

In this manuscript we have introduced a mechanism allowing for extremely long transients once a catastrophic bifurcation has occurred in a simple model for semiarid ecosystems (Kéfi *et al.* 2007). This mechanism, so-called ghost, arises in the vicinity of saddle-node bifurcations that leave, after the collision of a stable node with a saddle, a saddle-node remnant in phase space (Strogatz 2000). Saddle-node bifurcations are usually found in bistable systems. Evidences of bistability in drylands have been recently reported by means of global field studies (Berdugo *et al*. 2017).

Previous works suggested the ghost effect as a transient-generator mechanism in ecological systems (Sardanyés & Fontich 2010). Systems undergoing saddle-node bifurcations are typically bistable. This means that before the bifurcation occurs two stable fixed points are present and, depending on the initial conditions, one of them will be reached. For the semiarid ecosystems model one stable state is given by an ecosystem with vegetated area and available fertile-soil and another stable state involving a full desert area. Once the bifurcation occurs, the system becomes monostable and there exists a single fixed point in phase space, which is called to be globally stable. In this scenario, and when the bifurcation parameter is close to its bifurcation value, large enough initial conditions suffer a long delay due to the ghost. That is, although the system will become extinct, the transient towards extinction undergoes a very long plateau before the collapse. Under this situation, a system that will suffer a collapse may seem to be in its resilient phase, and anticipation for recovery would be extremely difficult.

The model analysed in this manuscript considers an ecosystem with three variables: vegetated soil, fertile soil, and desert. A saddle-node bifurcation and a saddle remnant have been identified at increasing the degradation rate of the soil. As mentioned above, the ghost effect and its associated inverse square-root scaling law have been identified. We must notice that the ghost effect is a local effect. It means that as the bifurcation parameter (*d* in the model analysed) increases beyond its critical value, the time to extinction largely decreases, and the ghost effect disappears. However, semiarid ecosystems poised near to the critical value could exhibit these dynamics, and the determination of stable, resilient state could be wrongly interpreted, having serious consequences.

Despite the clear difficulty of finding critical bifurcation values in real ecosystems, it has been recently discussed the possibility to identify ecological turning points by means of the so-called early warning signals (Carpenter & Brock 2006; Carpenter *et al.* 2012; Scheffer *et al.* 2015). In this sense, once warning signals may be detected, the delaying properties of the ghost could allow for strategies to prevent ecosystems’ collapses. In this manuscript we investigate the effect of external interventions in semiarid ecosystems close to a catastrophic shift. By means of small increases in the fraction of vegetated areas, ecosystems’ stability can be maintained since dynamical trajectories are constantly sent near the ghost, where time delays are extremely long.

We have quantified how these interventions (in terms of newly vegetated areas and frequency of application) might be useful depending on how far the system is placed beyond the bifurcation. Here it is important to highlight two aspects. The first one is that delayed transitions are extremely local phenomena. That is, the ghost effect rapidly disappears as *d* increases beyond *d*_*c*_. Another one is that our approach is neglecting spatial correlations, which are known to be crucial in ecological systems (Pascual 1995; Bascompte & Solé 1995; Hanski 2002). It is known that space usually enlarges transients (Hastings 2001; Hastings 2004). However, the impact of space in delayed transitions remains unknown for ecological systems as well as for other spatially-extended systems suffering saddle-node bifurcations.

Another important point is that the success of the interventions might depend on the underlying spatial patterns of the semiarid ecosystems. In this sense, several distinct spatial patterns have been characterized for semiarid ecosystems. Depending on the climatic conditions (i.e., moisture, slope, degradation) the vegetation can be homogeneous or organized in gaps, stripes and spots (von Hardenberg *et al.* 2001; K. Gowda, *et al*. 2014). Also, the so-called multi-scale patterning combining both environmental conditions and the action of territorial animals has been suggested to determine different types of self-organized spatial patterns in semiarid ecosystems (see e.g., Tarnita *et al*. 2017). That is, for the same fraction of vegetated area introduced during the interventions, differences in the dynamics could arise depending on where vegetation is re-planted and thus on the specific spatial patterns of the existing vegetation at the moment of applying the interventions. These points should be addressed in future research.

## Acknowledgments

The authors thank the members of the Complex Systems Lab for useful discussions. We also thank Sergi Valverde for facilitating the software used to build the spheres of Fig. 1, as well as Antoni Guillamon and Ernest Fontich for useful comments. We want to specially thank J. Tomás Lázaro for his technical assistance. This study was supported by Secretaria d’Universitats i Recerca del Departament d’Economia i Coneixement de la Generalitat de Catalunya, the Botin Foundation, by Banco Santander through its Santander Universities Global Division and by the Santa Fe Institute. JS has been partially funded by the CERCA Programme of the Generalitat de Catalunya. The research leading to these results has received funding from ”la Caixa” Foundation.

## Exploiting delayed transitions to sustain semiarid ecosystems after catastrophic shifts

### 1. FIXED POINTS AND STABILITY

The mathematical model we are studying is given by the next couple of differential equations:

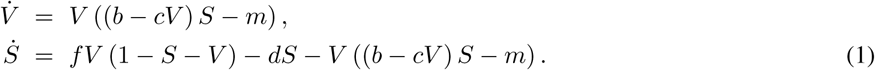

The fixed points are obtained setting 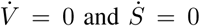 simultaneously. By doing so we can obtain a fixed point 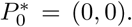. The (linear) stability of the fixed points is obtained from 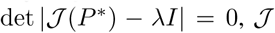 being the Jacobian matrix of Eqs. (1) and *I* de identity matrix. *λ* correspond to the eigenvalues, whose sign determine the stability. The Jacobian matrix reads:

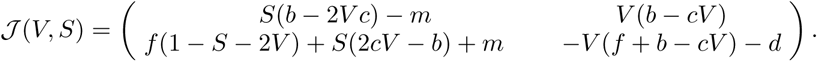

The stability of the fixed 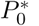 is easy to obtain and indicates that this equilibrium is locally stable, since both eigenvalues are negative 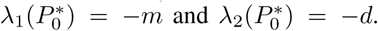 Notice that this point is locally stable, thus after the saddle-node bifurcation ocurring at *d* = *d*_*c*_.

The calculation of the other fixed points is rather cumbersome due to the structure of the model. The biologically-meaningful fixed points are found by means of the nullclines of (1), defined as the curves where either 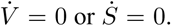 The nullclines are given by:

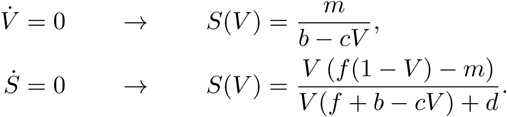

The intersections of these nullclines provide the equilibrium points of the system since 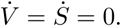 Figure S1 displays the shape of these nullclines and how they change in terms of parameter *d*. In Fig. S1a we set *d* = 0.1, and both nullclines intersect twice inside the phase space. This scenario corresponds to bistability since two stable fixed points 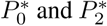 are found, separated by an unstable one 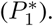. We notice that numerical solutions of Eqs. (1) as well as the pseudo-potential analysis indicate that the fixed point 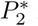 is stable (see Figs. 1 and 3 in the main manuscript). As *d* is increased towards its bifurcation value *d*_*c*_ both fixed points 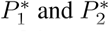 approach each other, as displayed in Fig. S1b. The value of *d*_*c*_ has been computed very accurately (see Section 2 below). At the bifurcation, both fixed points 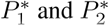 collide and disappear. For *d* > *d*_*c*_ the only fixed point is 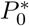 which is globally stable (see Fig. S1c).

**FIG. 1:**
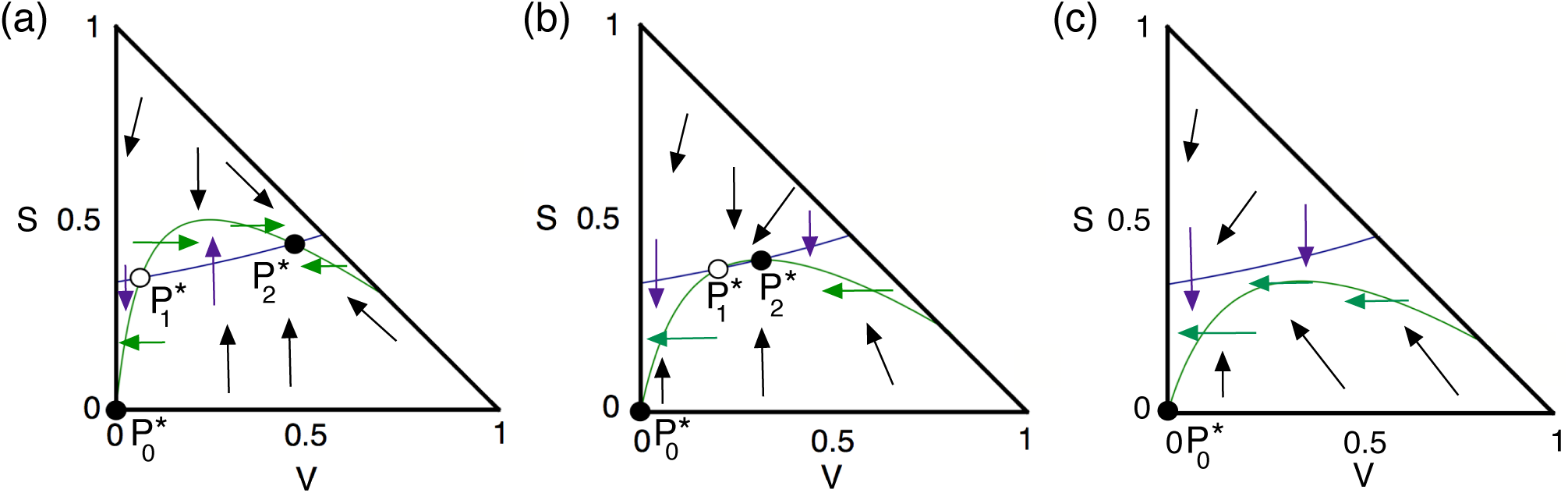
Nullclines obtained from Eq. (2) (violet line) and Eq. (3) (green line). The directions of the flows are indicated with small arrows. Here the intersections of the nullclines are fixed points (black: stable; white: unstable). Three different dynamical scenarios for several *d* values are displayed: (a) bistability with persistence of about 50% of a fraction of the vegetation (given by the stable fixed point 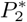), with *d* = 0.1 < *d*_*c*_; (b) decrease of the fraction of vegetation and approach of the unstable fixed point 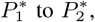, with *d* = 0.2104 ≲ *d_c_*. Finally, in (c) we set *d* = 0.32 > *d_c_*, and the fixed point involving the desert state 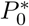 becomes globally stable since both fixed points 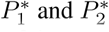 have coalesced. In all the panels we set *b* = 0.3, *c* = 0.15, *m* = 0.1, and *f* = 0.9.

#### 2. SHARP CALCULATION OF THE CRITICAL DEGRADATION RATE OF FERTILE SOIL

An accurate calculation of the bifurcation value of the degradation rate of fertile soil is a key point in our analyses since we are analyzing a phenomenon that occurs very near the saddle-node bifurcation. The bifurcation value can be usually calculated analytically from the expressions of the two fixed points involved in the saddle-node bifurcation: fixed points 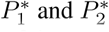 in our system. When the fixed points can not be calculated analytically, other strategies can be followed. One possible way of obtaining a very sharp calculation of this bifurcation value is by means of the so-called double discriminant theory. Computing bifurcation values with high accuracy is usually an extra difficulty in many problems, requiring an important numerical computation effort. However, when the vector field is of polynomial type (like in our model) or even rational, algebraic tools can help to find such values with more precision. The key point is to transform the original problem of looking for zeroes into a more simplified well-posed one.

This method is based on the so-called *discriminant* and *resultants* of one or a pair of polynomials. Roughly speaking, modulo some suitable constant, it is the product of the differences of the roots of the polynomial, counted with their multiplicity. It is a symmetric function of them. Closely related to this, we have the definition of *resultant* of two polynomials, Res (*p*(*x*), *q*(*x*)), defined as the product of the difference between the roots of *p* and the roots of *q* (counted again with their multiplicity). Therefore, Res (*p, q*) = 0 if and only if *p*(*x*) and *q*(*x*) have a common root. The discriminant Δ of a polynomial *p*(*x*) can be expressed as a constant multiplied by the Res (*p*(*x*), *p*'(*x*)). Zeroes of the resultant (or the discriminant) determine in many situations a change in the topology or in the behaviour of the problem (related to the change in the number of zeroes of *p*).

This method based on the computation of resultants can be used in many biological models when the vector field is polynomial. This is the case of our problem, where the system has an expression of the form

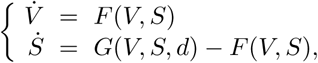

where *d* is taken as the parameter governing the bifurcation analysis, and

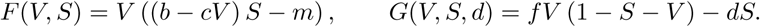

The amount and behaviour of equilibrium points is determined by the zeroes of the system

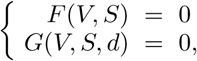

that is, the intersection points between both nullclines. Since the polynomials *F* and *G* depend on two variables, the method used is an extension of the presented above. It is based on the computation of the so-called *double resonant* (see, for instance Niu & Wang 2008; García-Saldaña *et al.* 2014; García-Saldaña & Gasull 2015; and Ferragut *et al.* 2016 and references therein). Since we are interested in the bifurcation value of the degradation rate of fertile soil, given by parameter *d*, we will fix all other model parameters at the same values used throughout all the analyses of our manuscript. Specifically, we are interested in *d*_*c*_ when *b* = 0.3, *c* = 0.15, *m* = 0.1, and *f* = 0.9. The computations to get *d_c_*, performed in `Maple`, consisted in the following steps:

**FIG. 2:**
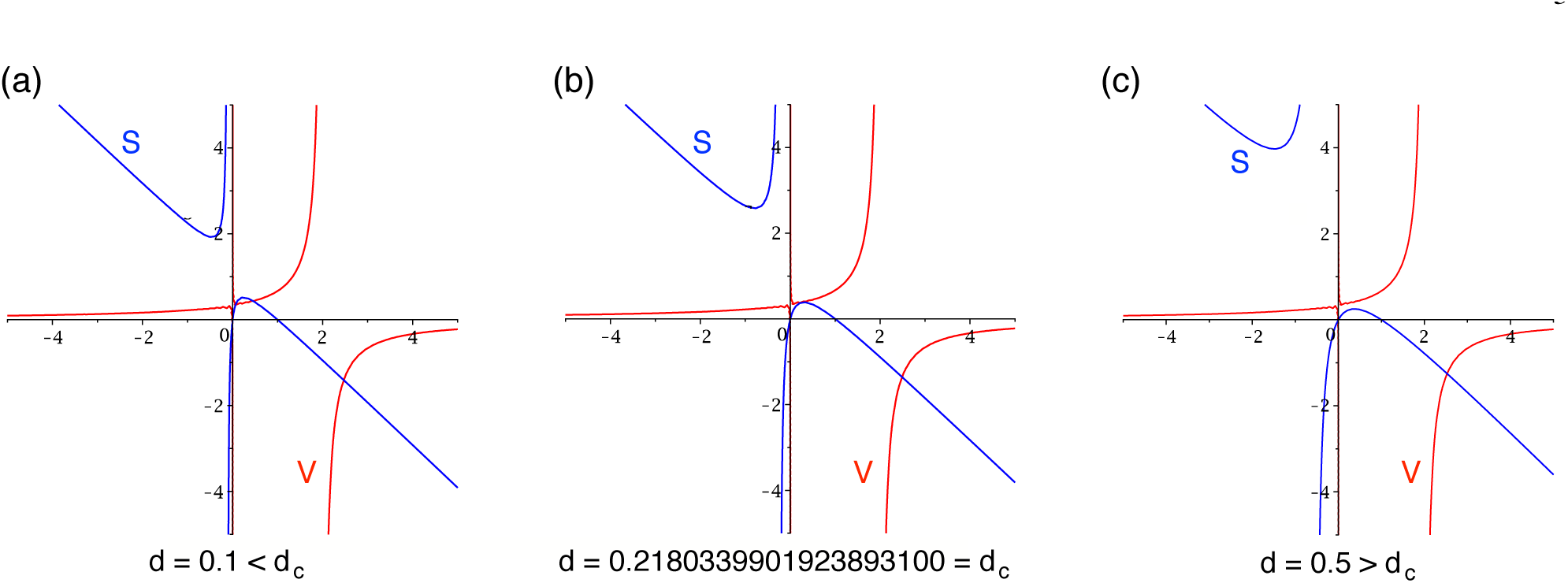
Shape of the curves *F* = 0 and *G* = 0 below (a); at (b); and above (c) the bifurcation value (see Section II). (a) The two curves intersect twice in the positive biologically-meaningful phase space below the bifurcation, here with *d* = 0.1 < *d_c_*. (b) The curves *F* and *G* are tangent just at the bifurcation value *d* = *d*_*c*_. (c) Above the bifurcation the two curves do not intersect each other, here with *d* = 0.5 > *d*_*c*_. In all of he panels we also set *b* = 0.3, *c* = 0.15, *m* = 0.1, and *f* = 0.9.

- Computation of

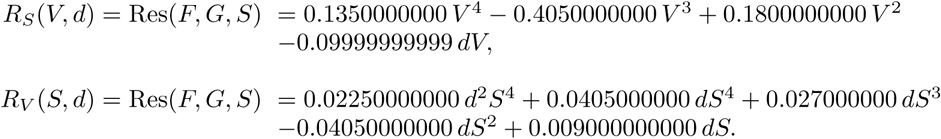
- We compute the resultants (the discriminant, modulo a constant):

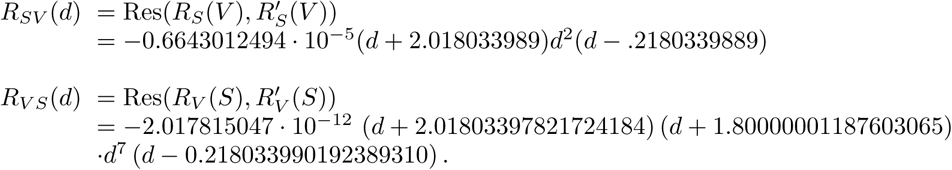
- Finally, the true (real) bifurcation values to consider are those common roots of this two last polynomials. In our case, since the paramater *d* must be non-negative, this is easily computed and provides the values *d*_1_ = 0 and *d*_2_ = 0.21803399019238931000 = *d*_*c*_.

It is straigthforward to check that for values *d* in *d*_1_ ≤ *d* < *d*_2_ there are three transversal intersections between the curves *F* = 0 and *G* = 0; at *d* = *d*_1_ just two (one of them tangential) and for *d* > *d*_1_ one transversal intersection (see Fig. S2, and movie1.tif in the online material for an animation about how the curves evolve at growing *d*). Remind that any intersection means an equilibrium point of the system. Notice that the previous method allowed us to obtain the bifurcation value *d*_*c*_ with a 20 decimal digits precision. This *d*_*c*_ value will be the one used in all of our analyses.

#### 3. NUMERICAL SOLUTIONS

The numerical solution for Eqs. (2)-(3) have been obtained using the fourth-order Runge-Kutta method with a step size Δ*t* = 0.1

#### 4. QUASI-POTENTIAL FUNCTIONS

The stability for many one-dimensional models can be derived from a so called potential function, *U*, connected to the dynamical flow through the expression

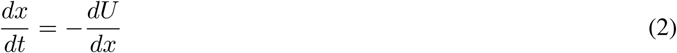

which gives a formal definition:

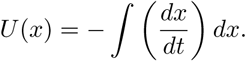

It can be shown that the minima and maxima of *U* correspond to the stable and unstable equilibrium points of the dynamics, respectively. This is in fact the formal expression of the (usually qualitative) pictures where a marble representing the state of the system rolls down from the bottom of one valley to another one as given parameters are changed. This is a well known picture for physical systems where forces (causing the dynamical trajectories) are derived from energy functions.

The mathematical definition of a potential function for more than one variable systems could be generalised as follows:

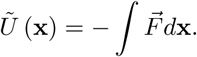

with x an n-dimensional vector state representing the population composition.

The potential energy 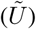 in point in the space is the energy needed to dissipate to achieve the minimum energy. This can have problems when we go to a system with more than one dimension. In order to maintain the same definition, the system have to be a conservative system. These systems are those that accomplishes that:

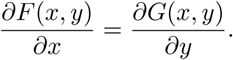

If this condition is true, the 2D potential function is given by:

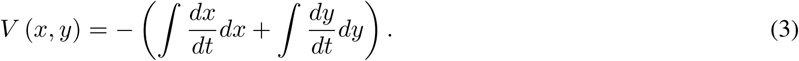

In our case, the system is not a conservative system. This is shown in the difference between the cross derivatives 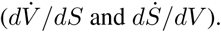.

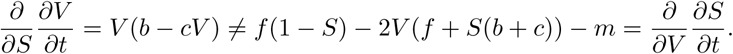

If the systems is not conservative, the potential function is not possible to define. For this reason, we define the quasi-potential function following the same equation (3), but it has not the physical potential meaning, but it can be used to illustrate the stability of the model. In our system, the integrals of the ODEs are:

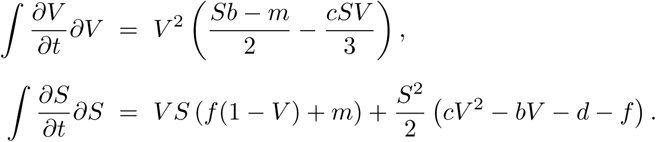

Then, the quasi-potential function is:

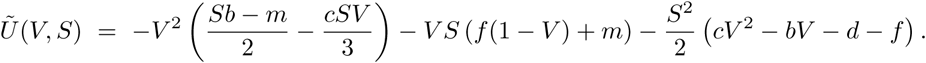

In the case that this integrals are not possible to compute or the underling equation system is not well defined, there are some articles that shows some tricks to perform an approximation to the define a quasi-potential function. This quasi-potential function is related with the numerical trajectories that can be computed by the numerical integration of the system of equations (Bhattacharya *et al.* (2011), Qiu *et al.* (2012), Li *et al.* (2013)). We will use the method implemented in Bhattacharya *et al*. (2011). This is based on the following expression:

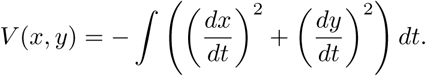

If we take a closer look to this formula, we can see that this formula is equivalent to the analytical definition.

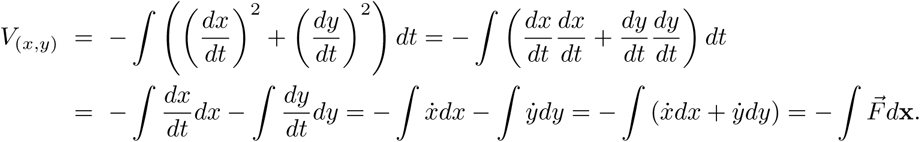

The numerical computation of the quasi-potential functions are done using the following algorithm.

**FIG. 3:**
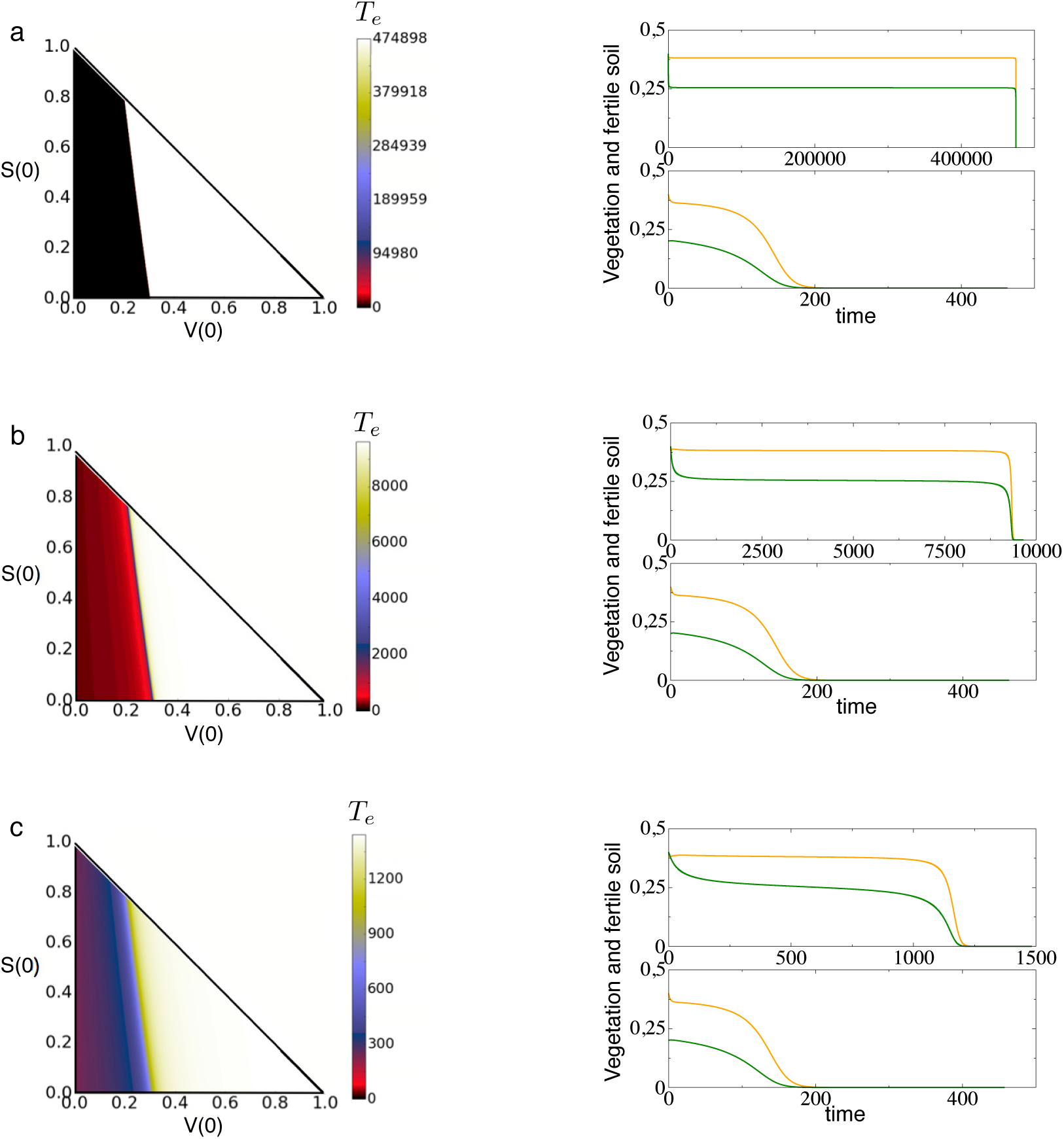
Extinction times, *T_e_*, computed numerically on the space of initial conditions (*V*(0), *S*(0)) setting *d* ≳ *d_c_*, with: (a) *d* = 0.218033995; (b) *d* = 0.21805; (c) *d* = 0.219. Below each panel we display transients for vegetation (green) and fertile soil (orange) states towards the desert state using two different initial conditions: (upper) the initial conditions close to the bifurcation collision point (*V*(0) = 0.4, *S*(0) = 0.4); and (lower) the initial conditions far away from the collision point (*V*(0) = 0.2, *S*(0) = 0.4).

1. Define a grid of initial conditions. In our case the mesh of initial conditions are from *S*_0_ = *V*_0_ = 0 to *S*_0_ = *V_0_* = 1 with Δ*S* = Δ*_V_* = 2 } 10^-3^. This defines a grid 500x500.
2. Compute the solutions of Eqs. (1) by numerical integration until the system reaches the equilibrium. In our case, we integrate Eqs. (1) using a 4-th order Runge-Kutta method with a time stepsize of δ_*t*_ = 0.1 for long times (we used *t* = 10^9^).
3. Compute the integral for each component.

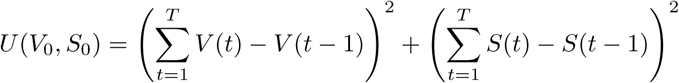

#### 5. NUMERICAL STUDY OF THE INTERVENTION STRATEGIES

In this Section we explain how the intervention method has been tested numerically. Two different processes have been considered: (i) increase of the fraction of vegetated region, labeled Δ*V*; and (ii) frequency of application of (i). To investigate the impact of the interventions in the system modeled by Eqs (2)-(3) in the main article, we have modified the fourth-order Runge-Kutta method to introduce these two processes. Below we display the algorithm (in pseudocode form) used to test computationally the designed intervention method.

##### Algorithm 1 Intervention algorithm using a 4th order Runge-Kutta method

set *V*_0_ = *V*(0) and *S*_0_ = *S*(0)

set *t* = 0

**for** *i in iterations* **do**

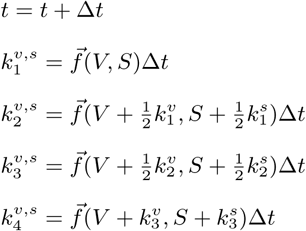

**if** *mod(t; freq*) == 0 **then**

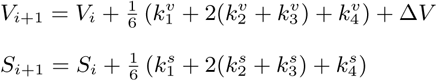

**else**

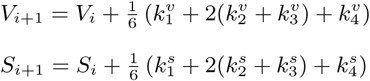

Here *V*_0_ and *S_0_* are the initial conditions; *t* is time (in arbitrary units); Δ*t* is the constant time step size; 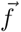 is the field (differential equations). The function *mod(t, freq*) = 0 indicates when the vegetation increment Δ*V* is introduced at a given frequency *freq*.

**FIG. 4:**
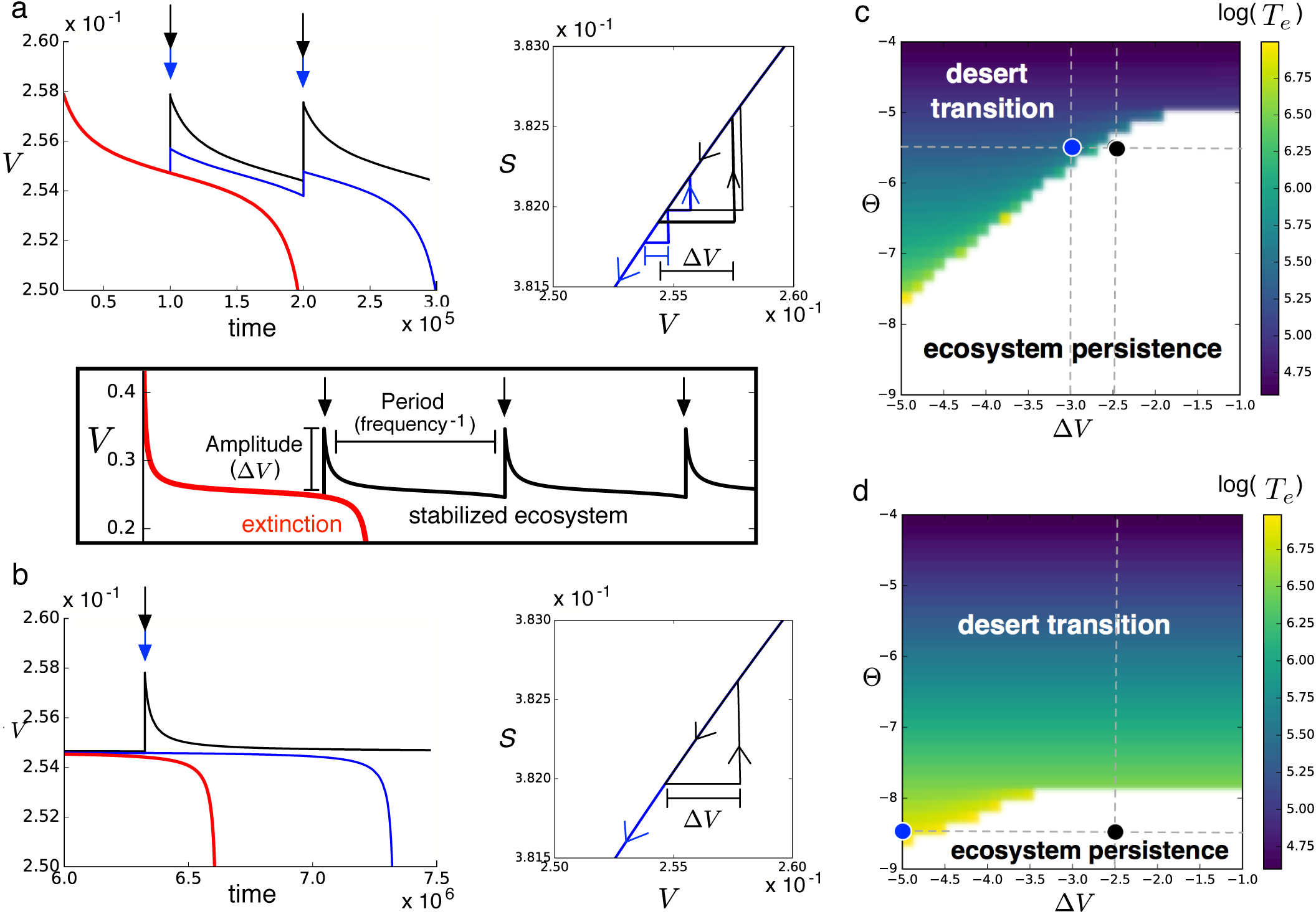
(a-b) Intervention strategies allowing to maintain ecosystems persistence after the bifurcation. The time series in panels a and b display three different dynamical outcomes beyond the bifurcation: the red trajectory indicates the extinction of the ecosystem without interventions. The blue line indicates an intervention that is not able to preserve the ecosystem; and the black time series corresponds to successful interventions able to preserve the vegetated state. Note that the intervention can be performed by adding a small fraction of vegetation (Δ*V*) at a given frequency (see black rectangle between panels a and b). The different dynamics can be visualized in the phase space (*V, S*) for each case. The two examples of panels a and b correspond to the dynamics obtained at a given distance of the bifurcation (parameter *θ*) and for a given value of Δ*V* in panels c and d, respectively.

---

**FIG. 5:**
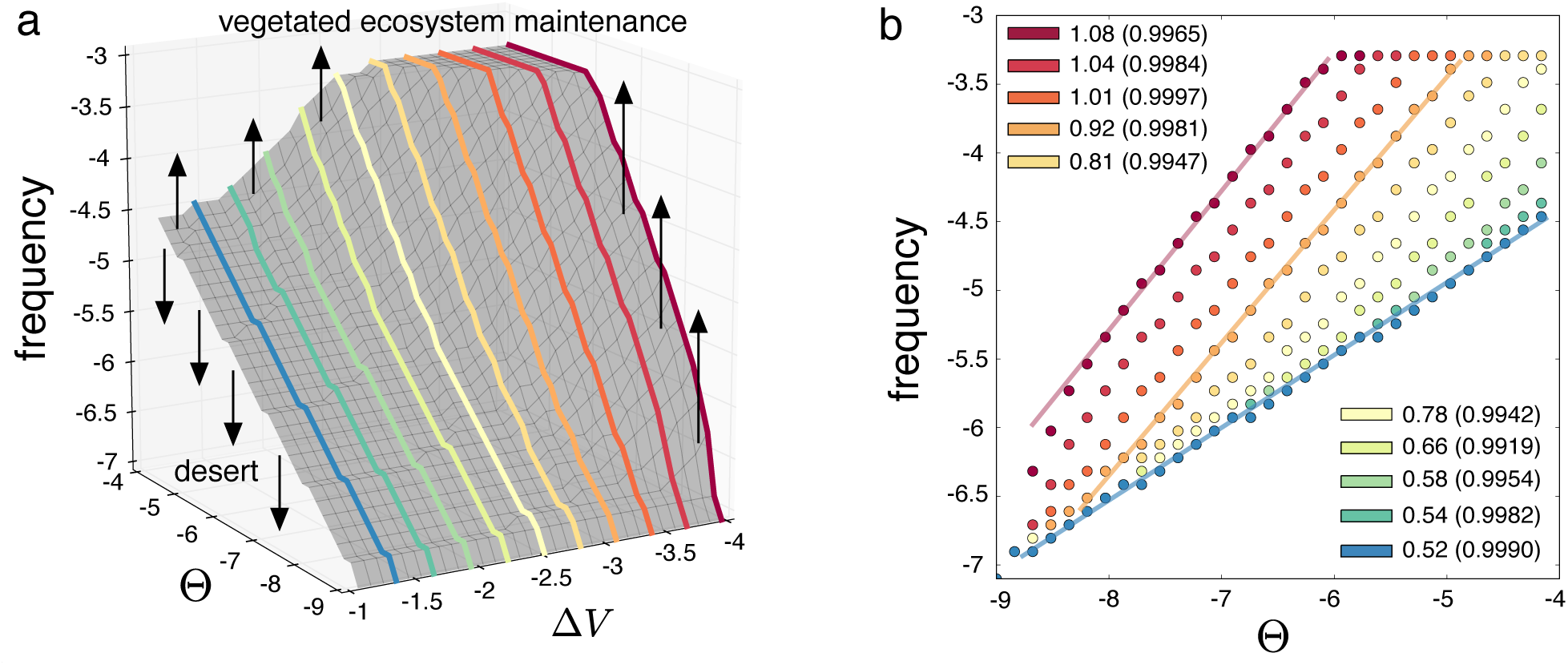
(a) Surface separating the regions for which the interventions allows to maintain the vegetation and the ones that fail to do so. The surface (a) is Pareto optimal: it is the minimum frequency and the minimum intervention magnitude needed for the maintenance of the vegetated ecosystem, (b) Cost increases in terms of frequency depending on Θ i.e., on the distance to the bifurcation value for different values of Δ*V*, displayed in panel a. Depending on the magnitude of the intervention, the frequency needed to hold the vegetation increases with different slopes as *d* increased beyond the bifurcation value. From red to blue, the frequencies of the interventions increases although the slope decreases. The straight lines in b display three examples of power-law fits. These fits have been performed for all of the cases represented in b. Inside the plot we display the scaling exponents of these power-law fittings and the values inside the parenthesis display the correlation coefficients of these fittings. The parameters used are the same used in the main text.

**FIG. 6:**
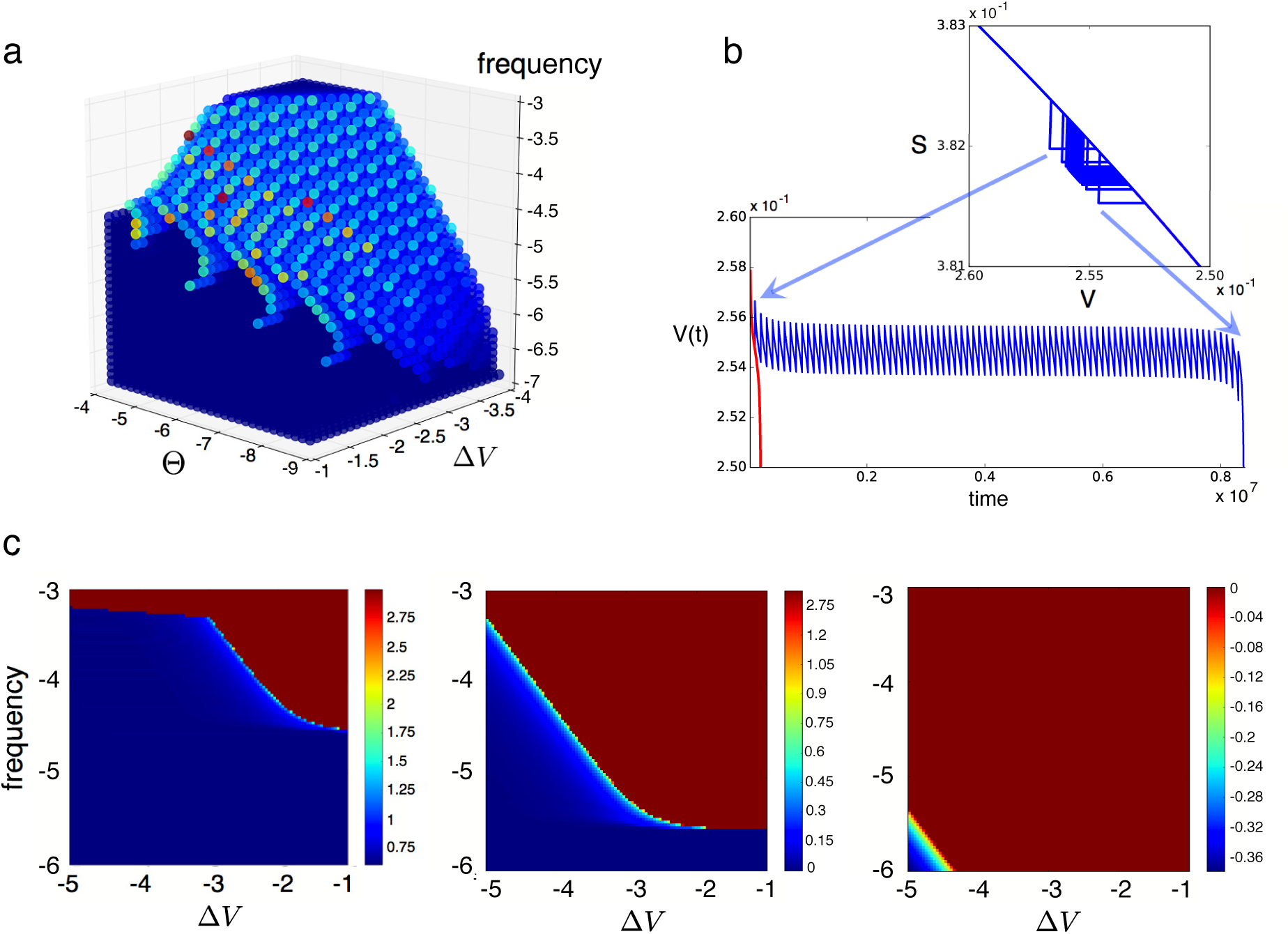
(a) Increase of the extinction times with respect to the non-intervention case in conditions of extinction. An example of elongation of survival time is illustrated in (b), where the intervention (*freq* = 10^-5^ and log_10_ Δ*V* = 2.7127) increases the time from 2 } 10^5^ (red) to 8.5 } 10^6^ (blue) at log_10_ Θ = −5.5. (c) Increase of extinction times, *T*_*e*_, depending on the intervention magnitude Δ*V* and frequency (sampling 100 } 100 points in the parameter space space (Δ*V, freq*)). From left to right : Θ = 10^−8^, Θ = 10^−6^, and Θ = 10^−4^. The other parameters are the same used for the previous figures.

